# Comparison of concentrations of lead (Pb) in meat from wild-shot common pheasants killed using shotgun pellets principally composed of lead, iron (Fe), bismuth (Bi) and zinc (Zn)

**DOI:** 10.1101/2023.05.12.540551

**Authors:** Rhys E. Green, Mark A. Taggart, Maider Guiu, Hayley Waller, Sabolc Pap, Rob Sheldon, Deborah J. Pain

**Affiliations:** Conservation Science Group, Department of Zoology, University of Cambridge, Downing Street, Cambridge CB2 3EJ, UK; RSPB Centre for Conservation Science, The Lodge, Sandy, Bedfordshire SG19 2DL, UK; Environmental Research Institute, University of the Highlands and Islands, Castle Street, Thurso KW14 7JD, U; 78 Riverdene Road, Ilford, Essex, IG1 2EA UK; School of Biological Sciences, University of East Anglia, Norwich Research Park, Norwich NR4 7TJ, UK

## Abstract

The source of almost all of the lead (Pb) found in meat from carcasses of wild-shot small game animals is often thought to be small embedded fragments of the lead shotgun pellets usually used by hunters to kill them. Available circumstantial evidence supports this conjecture, but an unknown proportion of the lead in game meat might be biologically-incorporated and absorbed by the game animal from ingested shotgun pellets and other environmental sources, such as soil and residues from mining. A critical test comparing lead concentrations in meat from animals known to have been killed using different types of shotgun ammunition has not been performed until now. We compared lead concentrations in samples of edible meat from carcasses of 27 wild-shot common pheasants (*Phasianus colchicus*) from which only lead shotgun pellets were recovered post mortem with concentrations for 20 birds from which only iron pellets were recovered. Shotgun pellets were removed from the meat samples before analysis. The mean concentration of lead was about 30 times greater in the meat of pheasants shot with lead than those shot with iron and was similar to mean concentrations of lead reported previously from Europe-wide samples of meat from wild-shot small game animals, including pheasants, killed using unknown types of ammunition. These results support the hypothesis that changing the type of shotgun ammunition in use for hunting from lead to iron would greatly reduce the concentration of lead in meat from wild-shot small game.

## 1. Introduction

Meat from free-ranging common pheasants (*Phasianus colchicus*) killed by hunters using shotguns is often sold and eaten by humans in the United Kingdom (UK) and elsewhere. In the UK, almost all pheasants are shot using shotgun pellets composed principally of lead (Pb) [1-4]. Lead shotgun pellets often fragment upon impact with the bodies of gamebirds, leaving small lead particles embedded in the meat that are difficult for consumers to detect and remove and from which a greater proportion of the lead is likely to be absorbed [1,5]. Absorbed lead impairs nervous, cardiac, renal, immune, endocrine and other functions in humans [6,7]. Since the introduction of regulations to reduce lead in, for example, petrol and paint, the primary route of exposure of humans to lead in the European Union (EU) and UK has become the diet [6]. Dietary lead derived from lead ammunition can contribute substantially to this exposure in people who consume game meat frequently [6,8]. The European Commission has set 0.1 ppm w.w. (wet weight) as the maximum level (EUML) permitted for lead in marketed meat (muscle) from domesticated animals, (Regulation EC1881/2006 - EC 2006), but no EUML has been set for lead in wild game meat.

There is clear evidence that meat from wild-shot small game animals contains substantial concentrations of lead [9]. This applies even when whole shotgun pellets have been removed from the meat sample prior to chemical analysis [1,9]. Replacement of lead shotgun ammunition by non-lead ammunition has been suggested as a potentially effective way to reduce dietary exposure of humans to ammunition-derived lead. In 2019, the European Chemicals Agency (ECHA) was asked by the European Commission to prepare a proposal to restrict the placing on the market and use of lead in ammunition under the EU Regulation on the Registration, Evaluation, Authorisation and Restriction of Chemicals (EU REACH). This would make hunting with lead shotgun ammunition and bullets unlawful in the European Union. The proposal may be implemented in 2024, if it is accepted and there are no delays to the regulatory process. Similar legislation is being considered by the UK Government under the UK REACH regulation, which has similar proposed timing.

Available evidence suggests that most of the lead in the meat of wild-shot small game animals killed with lead shotgun ammunition is likely to be from embedded lead fragments derived from the pellets used to kill the animal. This evidence includes X-ray studies, studies finding low lead concentrations in the meat of small game animals that are not lead poisoned and have not been shot, and studies of tissue lead concentrations in birds experimentally dosed with lead shot [9]. However, a proportion of the lead in game meat may be biologically-incorporated lead from other environmental sources. Biologically-incorporated lead accumulates in body tissues, especially the liver, kidney and bone, with lower concentrations usually being found in muscle, though lead concentrations in muscle may be elevated in lead-poisoned birds [10-12]. A study of stable isotopes of lead in the bones of wild red grouse (*Lagopus lagopus scoticus*) at three UK sites where the species is hunted by shooting (two in Scotland and one in England), indicated that the birds at both sites in Scotland had been exposed long-term to lead from shotgun ammunition by ingesting spent pellets. At the site in England, isotope ratios were consistent with a combined exposure to ingested lead gunshot and lead from past mining of a lead ore (galena) in the region [13]. Hence, whilst a lower concentration of biologically-incorporated lead is found in muscle than in bone, the potential exists for some of the lead in meat from wild-shot small game to be biologically-incorporated and derived from exposure to lead from sources other than embedded shot-in lead fragments.

Critical further evidence on this topic would be a comparison of lead concentrations in meat from small game animals between carcasses of animals killed using lead shotgun ammunition and those killed using non-lead shot. However, we are not aware of any robust studies which have made this comparison, although such information is available for large game animals killed with lead and non-lead bullets [14]. In this paper, we fill this evidence gap by comparing the concentration of lead in meat from free-ranging common pheasants known to have been killed using lead shot and three other types of shotgun ammunition.

## 2. Material and Methods

### 2.1. Compliance with Ethics Requirements

Carcasses of the dead common pheasants used in the study were all from birds killed legally by hunters in the UK and were obtained from food retailers and game meat wholesalers. The study did not involve any animal experiments.

### 2.2. Provenance of pheasant carcasses

We obtained 101 carcasses of free-ranging common pheasants, all of which had been shot on estates in England and processed and packaged for sale to consumers as ‘oven-ready’ prior to us acquiring them. Carcasses had been plucked and the head, neck, tarsi, feet and viscera had been removed. The carcasses were obtained as six batches, which differed as to source, date of acquisition and the co-author who processed the carcasses. Details of the batches are given in Supplementary Table S1. Batches 1 and 2 were provided to us for research purposes by a game supply business which intended to supply its customers with pheasant carcasses killed only using non-lead shotgun ammunition but was unsure about the types of ammunition actually being used. We purchased the other four batches from retailers of wild-shot pheasant carcasses without indicating to them that they would be used for research purposes.

### 2.3. Recovery of shotgun pellets

We unpacked and examined each carcass and attempted to find shotgun pellets by removing the skin and dissecting the muscles. We did not always attempt to recover all of the pellets which might have been present in each carcass and sometimes stopped searching when we had recovered one or two. Methods used to find shotgun pellets are described elsewhere [15]. We recovered at least one shotgun pellet by dissection from 54 carcasses. Pellets from each carcass were stored in a screw-topped polyethylene tube marked with a unique code.

### 2.4. Identification of the principal chemical element in shotgun pellets

When more than one pellet was recovered from a carcass, we performed qualitative tests to determine whether they were all of the same type or of more than one type. We determined surface colour, deformability/brittleness, attraction to a magnet and whether or not the pellet melted when touched with a hot soldering iron. These tests do not identify unequivocally the principal metallic element from which the pellet is composed, but they allow different types of pellets recovered from the same carcass to be distinguished. The methods and the metals they distinguish between are described elsewhere [2,3]. If the tests indicated that all pellets from the same carcass were of the same type, we selected one of the pellets at random for chemical analysis. If our tests found pellets of different types from the same carcass, we analysed one pellet of each type. We attempted to dissolve each of the pellets selected for chemical analysis in nitric acid. If the pellet did not dissolve in nitric acid alone, we used a 1:1 mixture of nitric and hydrochloric acids. We used an Inductively Coupled Plasma Optical Emission Spectrometer (ICP-OES; Agilent 5900 with SPS4 autosampler) to measure the concentrations of metals in the solution and to estimate from them the proportion of the mass of each pellet comprised of each metallic element. This method is described in more detail elsewhere [2-4]. We assigned each pellet to a principal metal type according to which metallic element comprised more than 50% of its mass.

### 2.5. Sampling of meat from pheasant carcasses

For Batches 1 and 3 one of us (REG) dissected off as much as possible of the edible meat from both breasts (pectoral muscles) and legs, diced it into approximately 1 cm cubes, mixed these thoroughly and took a sample of 30-50g of the mixture. A similar procedure was followed for Batches 2, 4 and 5, which were processed by DJP, but with only breast meat being sampled. For Batch 6 which was processed by RS, four samples, each of approximately 10g of meat were taken from each breast and leg, to give an overall sample of 30-40g. The samples were placed into individually-coded polythene bags, sealed and frozen before being sent for analysis.

### 2.6. Measurement of the concentration of lead in pheasant meat

We thawed the frozen samples of meat and examined them macroscopically to find and remove any whole shot not already detected and removed during dissection of the carcasses. This was aided by flattening the sample within the sample bag, then uplighting the sample with a large flatbed 44W LED work light. Any shot present were then silhouetted within a thin layer of translucent tissue. Samples were weighed, dried to constant mass, weighed again and milled to a fine powder. From each milled sample, 0.4g was digested in nitric acid and hydrogen peroxide and samples, certified reference material and blanks were analysed using an inductively coupled plasma optical emission spectrometer (ICP-OES; Agilent 5900). We expressed the concentration of lead in meat on a dry weight (d.w.) and wet weight (w.w.) basis. These concentrations and the limits of detection (LOD) for each analysis are shown in Supplementary Table S2.

### 2.7. Statistical Analysis

We assigned a value of half of the LOD to the three carcasses with lead concentrations in meat which were below the LOD. We then calculated arithmetic means of d.w. and w.w. lead concentrations in meat from carcasses from which we had recovered each type of shotgun pellet, after excluding results for two carcasses in which we found two different types of shot. We obtained 95% confidence intervals for the arithmetic means by calculating the 95% confidence interval of the mean of the log_e_-transformed values, back-transformed these limits and then applied a correction factor [16] to the lower and upper bounds to convert them to lower and upper confidence limits of the arithmetic mean.

Separately for carcasses of pheasants killed using lead shot and iron shot, we plotted cumulative distributions of the dry weight lead concentrations in meat and compared them by eye with the expected log-normal distributions based upon the mean and standard deviations of the log_e_-transformed values for each type. We used Kolmogorov-Smirnov one-sample tests to assess the statistical significance of deviations of the observed cumulative distributions from those expected from the fitted log-normal models [17].

We used Welch’s unequal variance *t*-test [18] to test the significance of differences between the means of the log_e_-transformed lead concentrations in meat from carcasses of pheasants killed using shotgun pellets principally composed of lead and of iron. We fitted ordinary least squares regression models to dry weight lead concentration data from carcasses of pheasants killed using shotgun pellets principally composed of lead and of iron. The dependent variable was the log_e_-transformed dry weight concentration of lead in the meat. Independent variables were shot metal type (binary variable: lead=1, iron=0) and batch code (a five-level factor). Although there were six batches (see Supplementary Table S1), the sixth batch only included pheasants killed using bismuth shot and therefore was not included. We fitted four models: (1) the null model with no main effects, (2) the main effect of metal type only, (3) the main effect of batch only and (4) the main effects of both metal type and batch. We calculated the small-sample version of the Akaike Information Criterion (AIC_c_) and AIC_c_ weights for each of the four models and the relative importance of the two variables [19]. We selected the model with the highest AIC_c_ weight. We calculated the relative importance of the two variables from the AIC_c_ weights of the models [19]. We used the regression coefficient of the model with the main effect of metal type only to estimate the log-ratio of the geometric mean concentration of lead in the meat of birds killed using lead shot to that for birds killed using iron shot. We back-transformed the 95% confidence limits of this log-ratio estimate by raising *e* to the power of each limit and then used a correction factor [16] to calculate the 95% confidence limits of the ratio.

## 3. Results

### 3.1. Types of shotgun pellets recovered

We recovered at least one shotgun pellet from 54 carcasses. Qualitative tests showed that the shotgun pellets recovered from 52 of the carcasses were of one type and that two carcasses had pellets of two types. We therefore conducted chemical analyses to identify the principal metallic element of 56 pellets. Of these, 28 (50%) were composed principally of lead, 22 (39%) were composed principally of iron and three each (5%) of bismuth and of zinc. The mean percentages by weight of each element in pellets assigned to each type were: lead – 94.9% (range 84.8 – 100.0%), iron – 96.7% (81.0 – 100.0%), bismuth – 98.5% (96.3 – 100.0%) and zinc 99.3% (97.7 – 100.0%). One carcass had at least one zinc and at least one iron pellet and another had both lead and iron pellets.

### 3.2. Concentration of lead in meat in relation to the metal type of shot used to kill the bird

We excluded results from the two carcasses with two different types of shot from our analyses of the concentration of lead in meat from carcasses of birds in relation to the types of shot used to kill them (Supplementary Table S2: #22 and #33). The arithmetic mean concentration of lead in meat was much higher for carcasses of pheasants from which lead shot were recovered than for those killed using iron, bismuth and zinc shot (Table 1). The arithmetic mean concentration of lead in meat from pheasants killed using lead shot was about 30 times higher than that for those killed using iron shot (Table 1). The difference between the means of log_e_-transformed concentrations for birds killed using lead and iron was highly statistically significant, both for dry weight (Welch’s *t*_*30*.*7*_ = 5.48, two-tailed *P* < 0.0001) and wet weight concentrations (Welch’s *t*_*31*.*3*_ = 5.30, two-tailed *P* < 0.0001). Inspection of Fig 1 suggests that the distributions of individual concentration values were approximately log-normal for both types of shot. The cumulative distributions for both shot types did not deviate significantly from those expected from the fitted log-normal model (Kolmogorov-Smirnov one-sample tests: lead, *D* = 0.113, *P* > 0.2; iron, *D* = 0.227, *P* > 0.2).

**Table 1.** Arithmetic mean concentrations of lead in samples of meat from wild-shot common pheasants in relation to the principal element of which a shotgun pellet recovered from each carcass was composed. 95% confidence intervals (C.I.) are shown for each parameter.

| Principal shot metal | *N* | Mean d.w. concentration (mg/kg) |  | Mean w.w. concentration (mg/kg) |  |
| --- | --- | --- | --- | --- | --- |
|  |  | Mean | 95% C.I. | Mean | 95% C.I. |
| Lead | 27 | 6.85 | 3.53 - 14.79 | 2.10 | 1.07 - 4.49 |
| Iron | 20 | 0.22 | 0.18 - 0.28 | 0.07 | 0.06 - 0.09 |
| Bismuth | 3 | 0.78 | - | 0.23 | - |
| Zinc | 2 | 0.16 | - | 0.05 | - |
| Ratio of means Lead:Iron |  | 30.5 | 12.7 - 64.5 | 28.9 | 12.0 – 61.4 |

**Fig 1.**
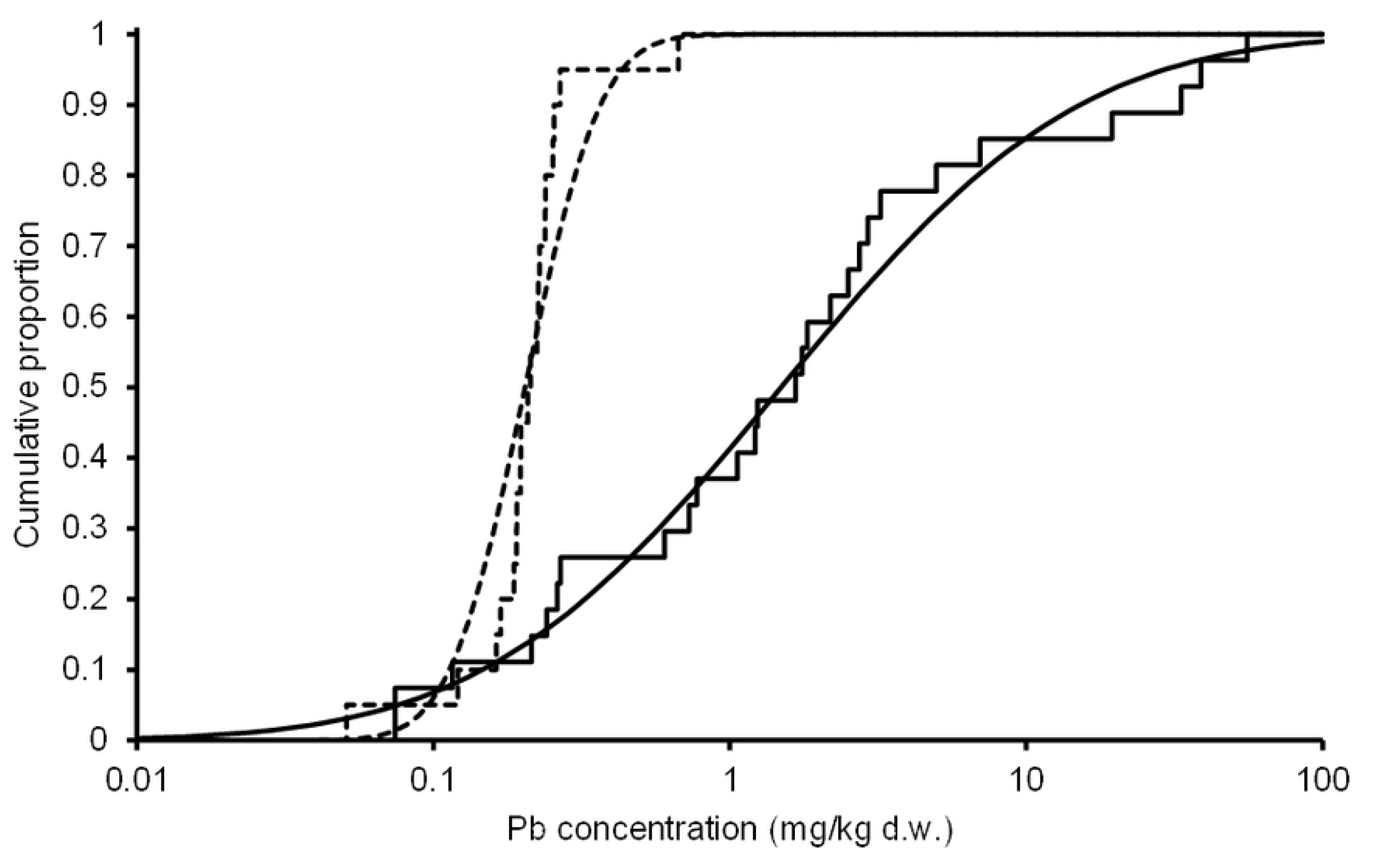
Cumulative distributions (stepped lines) of the dry weight concentration of lead in the meat from carcasses of wild-shot common pheasants from which a lead shotgun pellet (solid line) or an iron pellet (dashed line) was recovered. Curves show fitted log-normal distributions.

Our regression models of log_e_-transformed dry weight concentrations in meat were intended to assess the relative importance of differences between birds killed using lead (*n* = 27) and iron (*n* = 20) shot and among the five batches of carcasses. Comparison of AIC_c_ weights among the four main-effects models considered (see Statistical Analysis) showed that the model with only the effect of metal type (lead v. iron) had by far the highest AIC_c_ weight (0.955). The next highest weight value was 0.042 for the model with effects of both metal type and batch. The relative importance of the variable metal type was much higher (0.996) than for batch (0.045).

Our study included too few carcasses of birds killed using only bismuth and zinc shot (*n* = 3 and *n* = 2 respectively) to give reliable estimates of mean lead concentration and their confidence limits for these shot types. Meat from both of the carcasses of birds killed using zinc shot had low concentrations of lead (0.031 and 0.073 mg/kg w.w.). The mean lead concentration in meat from birds killed using bismuth shot was much lower than for those killed using lead shot (Table 1; individual lead values 0.069, 0.106 and 0.509 mg/kg w.w.). However, two of these values exceeded the EU Maximum Residue Level (0.1 mg/kg w.w.) for meat from domesticated mammals and poultry, which is a significantly higher proportion than for birds killed using iron shot (1/20 exceeded the threshold; Fisher exact test, two-tailed *P* = 0.034).

## 4. Discussion

Only 20 of the 54 pheasant carcasses with shot recovered which we used in our study were purchased from food retailers and game dealers (Tables S1 and S2). The other 63% had been requested by us from a supplier of game who was attempting to supply only birds killed using non-lead shot. Hence, our results should not be used to assess the proportions of wild-shot game killed using lead and non-lead shotgun ammunition.

The principal objective of our study was to compare the mean concentration of lead in meat from which embedded shot had been removed between carcasses of birds killed using different types of shotgun ammunition. Our most robust finding was that the arithmetic mean concentration of lead in meat from pheasants killed using lead shot was about 30 times greater than that from birds killed using iron shot. Regression analyses indicated no additional effect on lead concentration in meat of the batch from which the carcasses were obtained, which was included in the analyses because carcass batches almost certainly came from different locations in England and the three co-authors used slightly different methods when collecting meat samples for analysis. As far as we are aware, our study is the first to compare lead concentrations in meat from small game animals killed using lead and non-lead shotgun ammunition. A comparison of lead concentrations in meat from roe deer (*Capreolus capreol*us) and wild boar (*Sus scrofa*) killed using lead and non-lead (mostly copper) bullets showed that the concentration of lead was considerably higher in animals killed using lead than non-lead bullets for both species [14].

The arithmetic mean concentration of lead in the meat of pheasants known to have been killed using lead shot found in our study was similar to the mean derived from a large number of published studies reporting concentrations in meat from small game animals (including pheasants) from many locations in Europe, including the UK. The mean wet weight concentration for wild-shot small game animals sampled during the period 1991 – 2021 was 2.47 mg/kg w.w. [9], compared with 2.10 mg/kg w.w. for pheasants known to have been killed using lead shot in our study. The 95% confidence intervals of the two means overlap substantially. Our mean concentration for pheasants killed using lead shot is also similar to that from previously reported samples of UK pheasants [9]. The type of shot used to kill the small game animals whose data were used in the Europe-wide small game study [9] was unknown, but the similarity in mean lead concentrations is unsurprising, given that a high proportion of shotgun ammunition used for hunting in Europe is thought to be principally composed of lead.

The EU Maximum Residue Level (0.1 mg/kg w.w.) for meat from domesticated mammals and poultry was exceeded in 74% of carcasses of pheasants known to have been shot using lead shot (Supplementary Table S2). This proportion is broadly similar to that found in a Europe-wide small game study [9]. This proportion was much lower (5%) for carcasses of pheasants in our study known to have been killed using iron shot (Supplementary Table S2). None of the meat from pheasants killed using zinc shot had lead concentrations in the meat which exceeded the EUML. However, it was of concern that three birds were shot with zinc shotgun pellets (two only with zinc and one with zinc and iron shot) because zinc is toxic when ingested by waterbirds [20] and probably by other bird species and has not passed the US system for approval as a non-toxic shot type. It is clearly important that all alternatives to lead are non-toxic, rather than just non-lead.

Concentrations for two of the three carcasses of birds killed using bismuth shot exceeded the EU Maximum Residue Level, though only by a relatively small amount (largest value, 0.509 mg/kg w.w.). None of the 20 carcasses of pheasants killed using iron shot had a lead concentration as large as the largest value for the three birds killed using bismuth shot. It seems possible that the higher concentration of lead in this carcass (#54 in Table S2) might be due to the presence of an appreciable amount of lead, as a contaminant, in the principally bismuth shotgun pellets that killed it. However, the bismuth pellet analysed for this bird had only 0.02% of lead in it by weight, compared with <0.01% for the pellets recovered from the other two bismuth-shot birds, which had lower concentrations of lead in their meat. Because not all shot were recovered from all pheasant carcasses, it is also possible that the carcass of the bird containing bismuth shot with the highest lead concentration in its meat may also have been struck by one or more lead shotgun pellets which passed through the body or were embedded but not recovered, leaving behind residual fragments of lead. Whilst this is possible, the presence of more than one type of shot in the same carcass is rare [2-4].

## 5. Conclusions

The mean lead concentration for samples of meat from carcasses of wild-shot common pheasants from which only lead shotgun pellets were recovered were about 30 times greater than for birds from which only iron pellets were recovered. The mean lead concentration in meat from pheasants known to have been killed using lead shot was similar to mean concentrations of lead reported previously for many samples of meat from wild-shot small game animals killed using unknown types of ammunition. Mean lead concentrations in the meat of pheasants killed using bismuth and zinc shot were lower than for those shot using lead, but the sample sizes for these two shot types were too small to reach firm conclusions. Our results indicate that changing the type of shotgun ammunition in use for hunting from lead to iron would greatly reduce the concentration of lead in meat from wild-shot small game.

## Acknowledgments

We are grateful to Lincolnshire Game Ltd for assisting us with the supply of pheasant carcasses and to John Gregson (Waitrose & Partners) and Mark Avery for advice.

## Supporting Information Captions

**Supplementary Table S1**. Provenance of six batches of oven-ready carcasses of wild-shot common pheasants.

**Supplementary Table S2**. Dry weight (d.w.) and wet weight (w.w.) concentrations in samples of meat from 54 carcasses of wild-shot common pheasants.

